# Bat Accelerated Regions Identify a Bat Forelimb Specific Enhancer in the *HoxD* Locus

**DOI:** 10.1101/034017

**Authors:** Betty M. Booker, Tara Friedrich, Mandy K. Mason, Julia E. VanderMeer, Jingjing Zhao, Walter L. Eckalbar, Malcolm Logan, Nicola Illing, Katherine S. Pollard, Nadav Ahituv

**Affiliations:** Department of Bioengineering and Therapeutic Sciences, University of California San Francisco, San Francisco, CA, USA; Institute for Human Genetics, University of California San Francisco, San Francisco, CA, USA; Gladstone Institutes, San Francisco, CA, USA; Department of Molecular and Cell Biology, University of Cape Town, South Africa; Key Laboratory of Advanced Control and Optimization for Chemical Processes of the Ministry of Education, East China University of Science and Technology, Shanghai, China; Division of Biostatistics, University of California San Francisco, San Francisco, CA, USA; Division of Developmental Biology, MRC-National Institute for Medical Research, Mill Hill, London, United Kingdom; Randall Division of Cell and Molecular Biophysics, King’s College London, Guys Campus, London, United Kingdom

**Keywords:** Bat, Enhancers, *HoxD*, Limb Development, Evolution

## Abstract

The molecular events leading to the development of the bat wing remain largely unknown, and are thought to be caused, in part, by changes in gene expression during limb development. These expression changes could be instigated by variations in gene regulatory enhancers. Here, we used a comparative genomics approach to identify regions that evolved rapidly in the bat ancestor but are highly conserved in other vertebrates. We discovered 166 bat accelerated regions (BARs) that overlap H3K27ac and p300 ChIP-seq peaks in developing mouse limbs. Using a mouse enhancer assay, we show that five *Myotis lucifugus* BARs drive gene expression in the developing mouse limb, with the majority showing differential enhancer activity compared to the mouse orthologous BAR sequences. These include BAR116, which is located telomeric to the *HoxD* cluster and had robust forelimb expression for the *M. lucifugus* sequence and no activity for the mouse sequence at embryonic day 12.5. Developing limb expression analysis of *Hoxd10*-*Hoxd13* in *Miniopterus natalensis* bats showed a high-forelimb weak-hindlimb expression for *Hoxd10*-*Hoxd11*, similar to the expression trend observed for *M. lucifugus* BAR116 in mice, suggesting that it could be involved in the regulation of the bat *HoxD* complex. Combined, our results highlight novel regulatory regions that could be instrumental for the morphological differences leading to the development of the bat wing.

**Author Summary:** The limb is a classic example of vertebrate homology and is represented by a large range of morphological structures such as fins, legs and wings. The evolution of these structures could be driven by alterations in gene regulatory elements that have critical roles during development. To identify elements that may contribute to bat wing development, we characterized sequences that are conserved between vertebrates, but changed significantly in the bat lineage. We then overlapped these sequences with predicted developing limb enhancers as determined by ChIP-seq, finding 166 bat accelerated sequences (BARs). Five BARs that were tested for enhancer activity in mice all drove expression in the limb. Testing the mouse orthologous sequence showed that three had differences in their limb enhancer activity as compared to the bat sequence. Of these, BAR116 was of particular interest as it is located near the *HoxD* locus, an essential gene complex required for proper spatiotemporal patterning of the developing limb. The bat BAR116 sequence drove robust forelimb expression but the mouse BAR116 sequence did not show enhancer activity. These experiments correspond to analyses of *HoxD* gene expressions in developing bat limbs, which had strong forelimb versus weak hindlimb expression for *Hoxd10*-*11*. Combined, our studies highlight specific genomic regions that could be important in shaping the morphological differences that led to the development of the bat wing.

## Introduction

Vertebrate limbs show a wide range of morphological variety ranging from fins to limbs. The developing tetrapod limb is made up of three skeletal elements: the stylopod (humerus/femur), zeugopod (ulna/tibia, radius/fibula), and autopod (carpals/tarsals; metacarpals/metatarsals; phalanges) [1,2]. Autopods are highly specialized, composed of different numbers and lengths of digits, and exhibit varying degrees of interdigital soft tissue (webbing). Autopods are a hallmark of tetrapod diversity and are essential for adaptation to life on land, in the sea and in the air. Bats are an extreme example of this. To form a wing, bat forelimbs have gone through three major changes: elongation of digits II-V, retention of membranous tissue forming the inter-digital patagia (chiropatagium) and a relative reduction in the diameter of the ulna [3–5]. These morphological innovations are clearly apparent in bat fossils from 52.5 million years ago [6,7]. The genetic changes that led to the development of these specialized limb structures and mammalian flight are likely to have occurred prior to the radiation of the Chiroptera, one of the most diverse mammalian orders.

Enhancers can regulate spatiotemporal gene expression during vertebrate development [8] and nucleotide changes within them can lead to phenotypic differences, such as limb malformations [9]. For example, regulatory regions in the 5*’Hoxd* locus have been implicated in digit specification during mammalian autopod development and loss of interactions with these regions can result in limb phenotypes, similar to *Hoxd10*-*Hoxd13* deletions [10]. Nucleotide changes in enhancers have also been linked to morphological differences between species [11]. One such example is the *Prx1* limb enhancer. The replacement of the mouse sequence of this enhancer with the homologous bat *Prx1* sequence resulted in mice with longer forelimbs [12]. The recent availability of several bat genomes (*Myotis lucifugus, Myotis davidii, Pteropus vampyrus*, and *Pteropus alecto*) [13–15] now make it possible to identify specific nucleotide changes in the bat lineage, as compared to other mammals, that could have a role in the development of the unique limb morphology of the bat.

Various computational approaches have been used to identify regulatory elements that could be involved in species-specific morphological changes [16–20]. These include human accelerated regions (HARs) and human accelerated conserved noncoding sequences (HACNSs), which are highly conserved sequences that have acquired a disproportionate number of nucleotide substitutions since humans diverged from our common ancestor with chimpanzees [19,21,22]. Based on epigenetic marks, Capra and colleagues predicted that at least 30% of these noncoding HARs are developmental enhancers [23]. So far, 62 out of 92 tested HARs have shown enhancer activity in mouse transgenic assays, and 7 out of 26 HARs, where the activity of the human and chimp sequences were compared, showed differential enhancer activity [24]. These include the limb enhancer sequences HAR2/HACNS1, which showed no limb specific activity for the non-human homologous sequence [21], and 2xHAR.114, which displayed restricted limb activity for the human sequence compared to the chimpanzee sequence [23]. These findings indicate that the identification of accelerated regions could serve to detect sequences that function as gene regulatory elements and could possibly give rise to characteristic phenotypes among species.

Here, we set out to identify enhancers whose alteration could have contributed to bat wing development. We utilized mouse limb-specific ChIP-seq datasets and available bat genomes to identify bat accelerated regions (BARs). We identified 166 BARs with potential enhancer activity in developing limbs and functionally analyzed five of them in mouse, finding all five *M. lucifugus* cloned BARs (BAR2, BAR4, BAR61, BAR97, BAR116) to be functional limb enhancers. Comparison of the enhancer activity of mouse and *M. lucifugus* orthologous BAR sequences revealed expression differences for three of the four tested sequences (BAR4, BAR97, and BAR116), suggesting that these sequences could be accelerated in bats due to functional differences. Amongst them, *M. lucifugus* BAR116, which resides in a gene desert on the telomeric side of the *HoxD* locus, showed robust forelimb and weak hindlimb expression, a trend similar to bat *Hoxd10* and *Hoxd11* gene expression as we determined using whole-mount *in situ* hybridization on bat and mouse embryos.

## Results

### Computational analysis identifies 166 BARs

We sought to identify specific sequences that could be responsible for bat wing development. To generate a high-confidence list of candidate enhancers, we implemented a comparative genomics approach (Fig 1) that pinpoints bat accelerated regions (BARs), which are genomic sequences that are evolving very slowly in vertebrates, but experienced rapid sequence changes in the common ancestor of extant bats. We analyzed multiple sequence alignments of 58 vertebrates, excluding bat genomes (see methods; Fig 1), to generate 2.7 million vertebrate conserved sequences using PhastCons [25]. To focus our analysis on potential limb developmental enhancers, we constrained our search to conserved sequences that overlap with 39,260 ChIP-seq peaks for H3K27ac and p300 from embryonic day (E) 10.5 and E11.5 mouse limbs. These include two previously reported datasets [17,26] and an H3K27ac E11.5 developing mouse limb autopod dataset generated for this project (see methods, Fig 1). We then tested these candidates for statistically significant numbers of substitutions in the ancestor of four bats with sequenced genomes, compared to the set of vertebrate conserved sequences, using PhyloP [27] (see methods). Using four bats and numerous vertebrate genomes in our analysis assisted in reducing false positives that can result from sequencing, assembly, and alignment errors.

**Fig 1.**
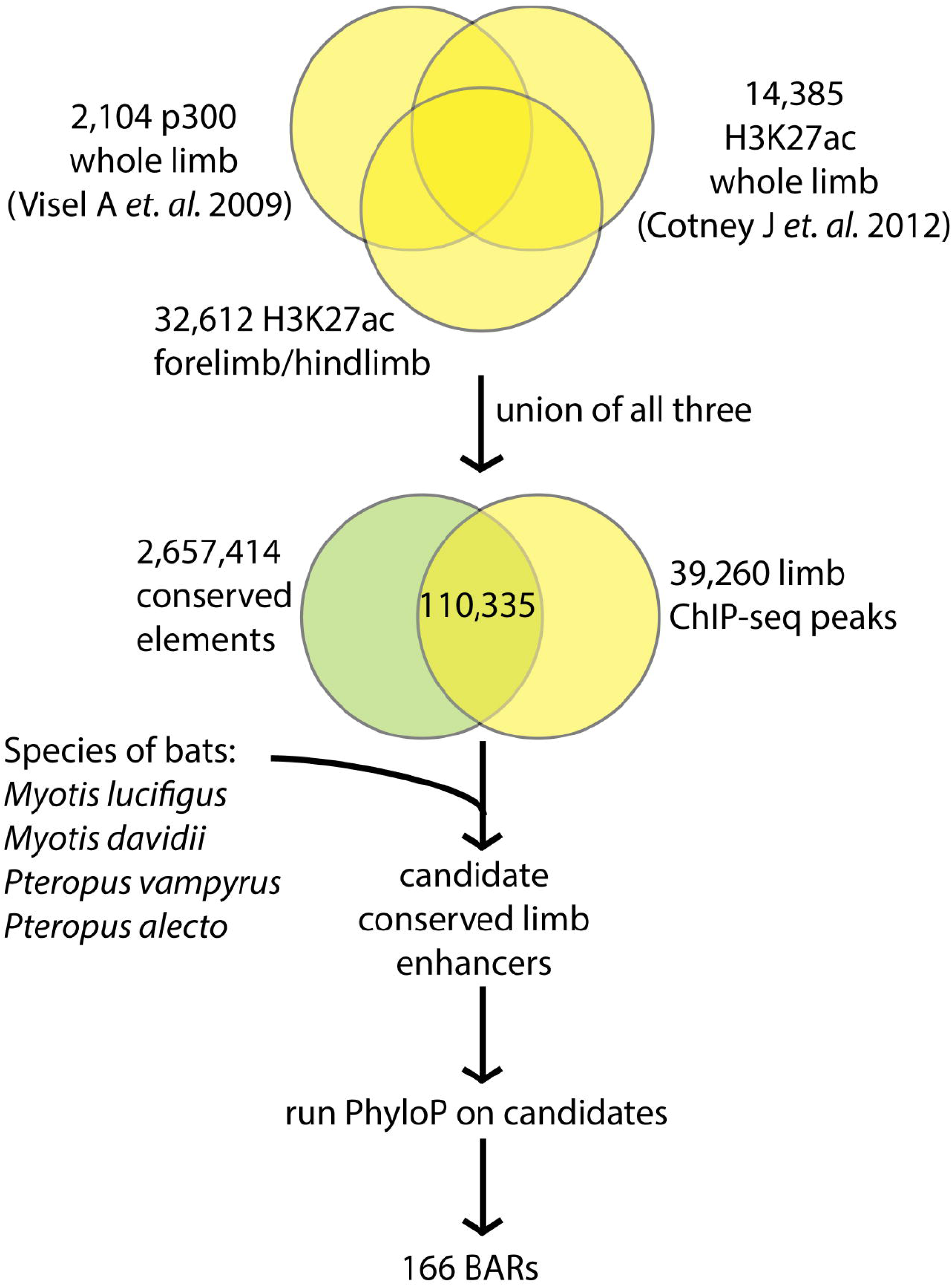
Computational pipeline to identify bat accelerated regions. Limb ChIP-seq peaks were unified, then overlapped with conserved regions and then scored with PhyloP values (0 to 20) by comparing *Myotis lucifugus, Pteropus vampyrus, Myotis davidii*, and *Pteropus alecto* to 48 available vertebrate genomes. A total of 166 BAR elements were identified as accelerated regions in bats (false discovery rate (FDR) < 0.05).

We identified 166 BARs that overlap genomic marks of active enhancers in developing mouse limbs and show significant evidence of accelerated substitution rates in the genome of the common ancestor of extant bats (false discovery rate (FDR) < 5%; S1 Table). Like many known developmental enhancers, the average BAR is 1,542 base pairs (bp) in length and does not overlap gene transcription start sites (TSS). We found that 73% of BARs are more than 20 kilobases (kb) from the closest TSS, 45% are more than 100 kb from a TSS, and five are in gene deserts that are greater than 1 megabase (Mb) across. Thirty-eight BARs are adjacent to transcription factors (TFs) involved in limb development (see methods; S1 Table) and overall we observed an enrichment for limb TFs near BARs (OR = 2.88, p-value < 0.0001; S2 Table). BARs were also found to cluster around each other more densely than expected (p<0.001; permutation test compared to the set of candidate limb enhancers from the various ChIP-seq datasets), with several clusters being adjacent to developmental genes. For example, we found five BARs (BAR4, 18, 22, 71 and 148) clustered near sprouty homolog 1 (*Spry1*), a gene associated with skeletal myogenesis [28,29]. In addition, for 4 out of 5 BARs (BAR4, BAR18, BAR22, BAR71, BAR148) near *Spry1*, we found more pleiomorphic adenoma gene 1 (*Plag1*) motifs in the inferred sequence of the bat ancestor than in the orthologous mouse sequence (all p-values<0.1; see methods). *Plag1* is a zinc finger protein whose loss in mice results in retarded growth [30] and it has been associated with stature in bovines [31], limb bone length in pigs [32] and height in humans [33].

### *M. lucifugus* and mouse BAR sequences differ in their transcription factor binding sites

We next set out to identify transcription factor binding site (TFBS) changes in each of the 166 BARs by estimating the sequence of the common ancestor of the four bat genomes (*M. lucifugus, P. vampyrus, M. davidii and P. alecto;* see methods) and comparing this ancestral bat sequence to the orthologous mouse sequence. We predicted TFBS in the mouse and ancestral bat sequences of each BAR and tested for significant loss or gain of TFBS of 745 TFs expressed in the developing limb using motifDiverge [34]. Most TFs only had significant changes in TFBS for a single BAR, but several showed consistent patterns of loss or gain across multiple BARs. When all BARs are analyzed collectively as a single sequence, 34 TFs have significantly more TFBS in the bat ancestor compared to mouse, and 146 TFs have significantly fewer TFBS (FDR<0.05, S2 Table).

The most striking TFBS changes in the ancestral bat BAR sequences were gains of sites for *Nr2c2*, *Sp4, Zfp281*, and *Zfp740* each of which is enriched in twelve or more BARs*. Nr2c2*, also known as the testicular nuclear receptor 4 (*Tr4*), is involved in osteoblast maintenance and differentiation [35,36]. Mice lacking *Tr4* do not have apparent skeletal abnormalities, however, they display a reduction in bone mineral density and long bone volume, showing premature aging, spinal curvature [37], and osteoporosis [35]. *Zfp281* and *Zfp740* are expressed in the developing limb [38] but have yet to be characterized for their limb function. Two additional TFBS gains are worth noting, *Egr1* and *Zic2/3*. The *Egr* genes are C2H2-type zinc finger proteins that function as transcriptional regulators with an important role in mitogenesis and differentiation. Specifically, *Egr1* is involved in mouse wound repair, endochondral bone repair and data suggests that *EGR1* is upregulated during skeletal muscle wound healing [39,40]. *Zic2* and *Zic3* belong to the C2H2-family of Zinc fingers, are known to be involved in morphogenesis and patterning during development and are associated with muscle and skeletal defects [41–44].

We also observed a significant depletion for specific TFBS when comparing the ancestral bat sequences to mice collectively over all BARs (S2 Table). By rank, the most depleted and fourth most depleted TFs were *OSR2* and *OSR1* respectively. Odd-skipped related genes, *Osr1* and *Osr2*, belong to the C2H2 Zinc finger family [45,46] and are expressed in the embryonic limb mesenchyme [47,48]. Both *Osr1* and *Osr2* are associated with osteoblast regulation, chondrogenesis [49,50], synovial joint formation, and their removal in mice leads to fusion of these joints [51]. Also worth mentioning are *Tgif1* and *Meis1*. *Tgif1*, the Thymine/Guanine interacting factor 1, is a repressor of TGF-β/Smad signaling, and is expressed in the developing limb mesenchyme [52]. *Meis1*, a TALE homeobox TF, is a marker of the stylopod region and its overexpression abolishes distal limb structures during development [53]. Combined, our results identify TFBS gains and losses in BARs that might have a functional role.

### *M. lucifugus* BARs are functional limb enhancers

To determine whether BARs are functional limb enhancers, we selected five BARs (BAR2, BAR4, BAR61, BAR97 and BAR116) and tested them for enhancer activity using a mouse transgenic assay. The BAR candidates were chosen based on their location, residing within 1Mb of a known limb developmental genes whose alteration leads to a skeletal or limb phenotype (Table 1; Fig 2). BAR2 resides near *Twist2*, a bHLH transcription factor which has been shown to terminate the Shh/Grem1/Fgf autoregulatory loop by repression of Grem1 expression in early limb morphogenesis [54]. BAR4 is in close proximity to *Spry1*, a known antagonist of FGF signaling [55], that along with other Sprouty proteins (*Spry2*, *Spry4*), is expressed in skeletal muscle stem cells, chondrocytes, limb buds, muscles and tendons during development [28,56,57]. BAR61 overlapped with the zone of polarizing activity (ZPA) regulatory sequence (ZRS), a previously characterized limb enhancer of *Shh* that when mutated leads to limb malformations in humans, mice, dogs and cats [58,59] (Table 1). BAR97 is in close proximity to *Spg20*, a gene that is expressed in the limb, face and brain during morphogenesis and is an inhibitor of BMP signaling that is linked to short stature and spastic paraplegias [60]. BAR116 is located on the telomeric side of the *HoxD* cluster (Fig 2) which is known to be an important regulator of skeletal development [61].

**Table 1.** BARs selected for mouse enhancer assays. BARs that were selected for enhancer assays, the limb-associated genes nearby, the limb phenotype caused by mutations in these genes and their references.

| BAR ID | Nearby Limb Genes | Limb-Associate Phenotypes (MGI, OMIM) & Tissue Expression | References |
| --- | --- | --- | --- |
| 2 | *Twist2* | Skeletal and muscle abnormalities | [93] |
| 4 | *Spry1* | Chondrodysplasia, Muscles, Tendons | [28,56] |
| 61 | *Shh* | Limb malformations | [58] |
| 97 | *Spg20* | Spastic paraplegias | [60] |
| 116 | *HoxD cluster* | Skeletal defects | [61] |

**Fig 2.**
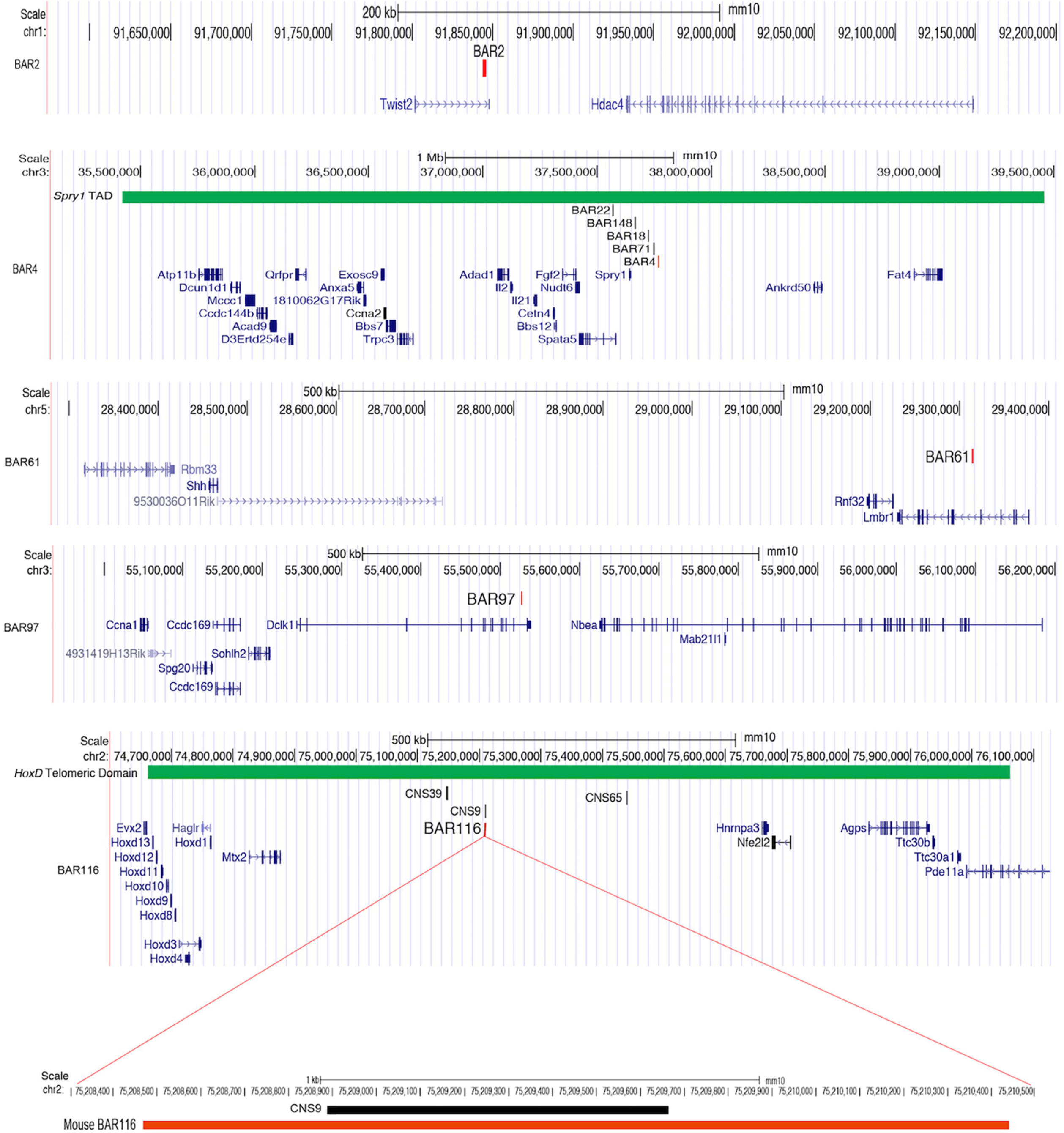
UCSC Genome Browser snapshots showing the location of each mouse BAR (BAR2, BAR4, BAR61, BAR97, BAR116). The BAR4 browser snapshot displays the ‘TAD Domain’ track (in green) containing a cluster of five *Spry1* BARs [69]. The bottom panel displays the mouse *HoxD* locus and a zoom-in of the overlapping regions between mouse BAR116 (~1.9kb in length) and mouse CNS9 (~700 bp in length) [74]. Both sequences were negative for limb enhancer activity in mouse assays. The *HoxD* ‘Telomeric Domain’ track is shown in green.

Regions spanning each of the five BAR candidate enhancers (Table 1; S1 Table) were amplified from *M. lucifugus*, cloned into the Hsp68-LacZ vector that contains an *Hsp68* minimal promoter followed by the LacZ reporter gene [62], and injected into single-cell mouse embryos. Transgenic embryos were harvested at E12.5. This stage was chosen since it is equivalent to CS16E in *Carollia perspicallata* and *Miniopterus natalensis* bat embryos, a stage when digits are identifiable and forelimbs (FL) lose their symmetry in the anterior to posterior (AP) axis compared to hindlimbs (HL) [63–65]. All assayed *M. lucifugus* BAR sequences showed limb enhancer activity in our transgenic mouse assay (Fig 3). *M. lucifugus* BAR2 was active in the limb autopod (3/5 embryos) but also demonstrated enhancer expression in the brain and nonspecifically throughout the whole embryo (4/5 embryos; S1 Fig). *M. lucifugus* BAR4 was positive for enhancer activity in the proximal limb (3/4 embryos; Figs. 3 and 4). *M. lucifugus* BAR61 demonstrated strong activity in the posterior-half of the autopod in FL and HL tissues (3/4 embryos) (Figs. 3 and 4). BAR97 displayed weak and diffuse enhancer expression in the posterior-half of the autopod (3/4 embryos; Figs. 3 and 4). *M. lucifugus* BAR116 showed strong enhancer activity throughout the proximal and distal FL, covering the entire autopod and zeugopod regions. A weaker enhancer activity in the proximal portion of the HL was also observed for BAR116 (3/5 embryos; Figs. 3 and 4). In total, all five examined *M. lucifugus* BAR sequences displayed enhancer activity in the developing forelimb or hindlimb.

**Fig 3.**
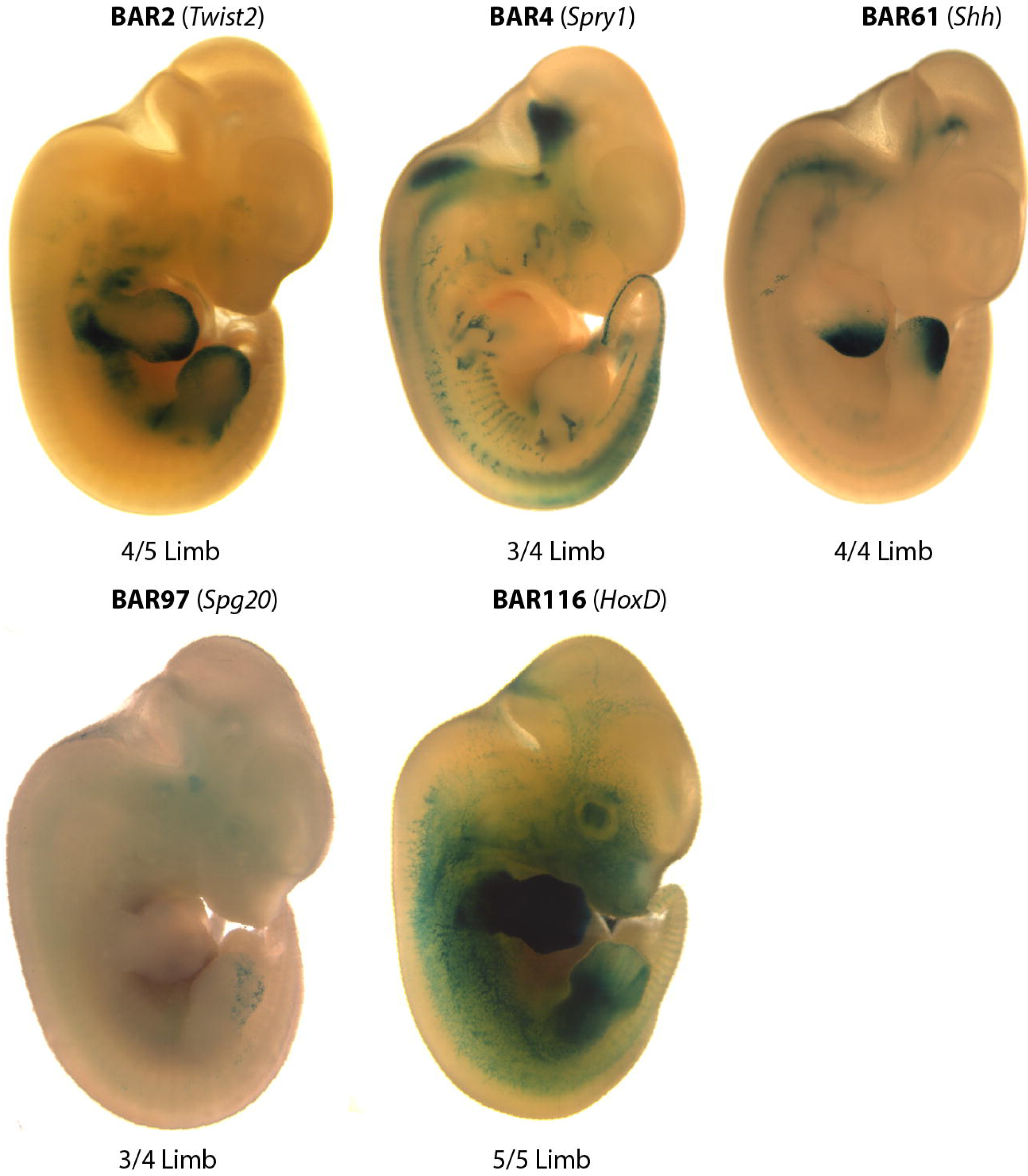
*M. lucifugus* BARs are active enhancers in the developing mouse limb. A representative mouse embryo (E12.5) showing limb enhancer expression pattern for each *M. lucifugus* BAR. Nearby limb-associated gene names are written in parenthesis with the number of embryos showing a limb expression pattern given below. *M. lucifugus* BARs were scored by the number of transgenic LacZ positive limb/ LacZ positive expressing embryos. All five *M. lucifugus* BARs have LacZ expression in the limbs.

**Fig 4.**
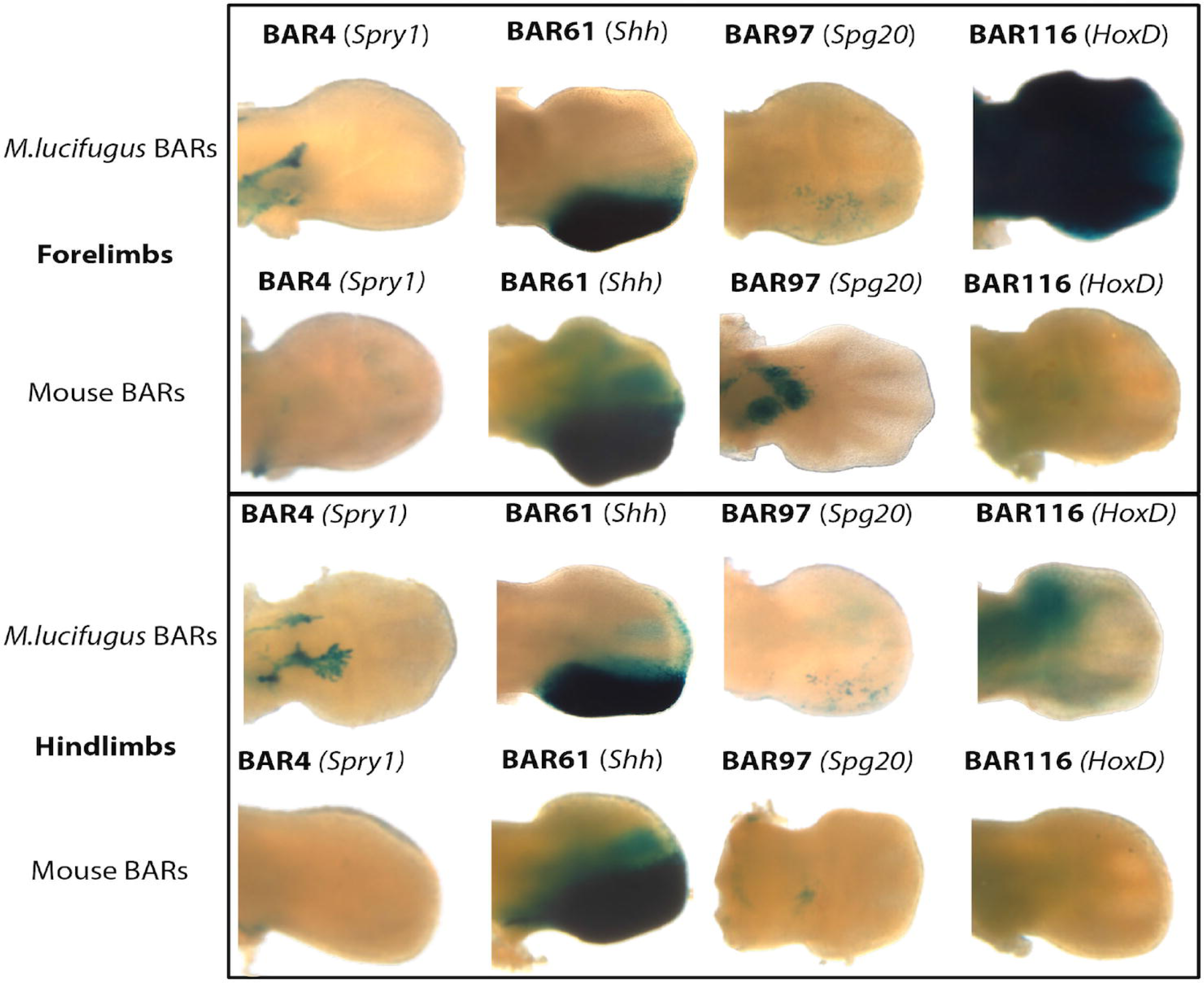
Comparison of enhancer expression patterns for bat and mouse sequences in forelimb and hindlimb. Representative mouse (E12.5) forelimbs (FLs) and hindlimbs (HLs) showing both *M. lucifugus* BAR and mouse BAR expression pattern. Three *M. lucifugus* BAR sequences (BAR4, 97, and 116) show differences in expression patterns as compared to the mouse BAR sequence. BAR61 (*Shh*) retains a similar expression pattern for both the bat and the mouse BAR sequences. Nearby limb-associated gene names are written in parenthesis next to the BAR ID.

### Orthologous mouse sequences demonstrate a divergent enhancer expression pattern

To compare the species-specific enhancer activity of our predicted BARs, we set out to analyze the orthologous mouse sequences of four BARs (BAR4, BAR61, BAR97, BAR116; S1 Table). Due to the nonspecific expression pattern of *M. lucifugus* BAR2, the orthologous mouse sequence was not analyzed. Regions covering each of the mouse BAR sequences were cloned into the Hsp68-LacZ vector and tested for enhancer activity at E12.5. Mouse BAR61/ZRS (*Shh*), was active in the posterior-half of the developing autopod (2/4 embryos, Fig 4; S2 Fig), similar to the corresponding *M. lucifugus* sequence (BAR61). However, the three other tested mouse BAR sequences (BAR4, BAR97, BAR116) showed differential enhancer activity. Mouse BAR116 (*HoxD*) and mouse BAR4 (*Spry1*) sequences were both negative for enhancer activity (Fig 4; S2 Fig). For mouse BAR4 it is worth noting that only one of the six positive embryos showed weak staining that was similar to *M. lucifugus* BAR4 and for mouse BAR116 none of the embryos showed similar enhancer activity, with the majority (3/4) being completely negative for LacZ (S2 Fig). Mouse BAR97 (*Spg20*) showed differential enhancer activity compared to *M. lucifugus* BAR97, being active in the midbrain, the zeugopod (2/4 embryos) and the forebrain (1/4 embryos, Fig 4; S2 Fig). Combined, our results suggest that the accelerated sequence changes observed in BARs could lead to differences in limb enhancer expression.

### The bat BAR116 sequence is essential for limb enhancer activity

Our injected BAR sequences included the PhastCons conserved sequences that defined the BAR element along with the flanking sequence under the ChIP-seq peak (S1 Table). We next wanted to determine whether the *M. lucifugus* BAR116 PhastCons sequence, rather than sequence differences in the flanking regions, is essential for the observed E12.5 limb enhancer activity. We generated a synthetic construct that has the mouse BAR116 PhastCons sequence along with the flanking *M. lucifugus* sequence. This bat-mouse BAR116 composite sequence was analyzed using a similar mouse transgenic enhancer assay at E12.5 and displayed inconsistent limb expression in two out of the four LacZ positive transgenic embryos (Fig 5, S2 Fig). The loss of limb enhancer activity that we observed for the bat-mouse BAR116 composite construct suggests that the *M. lucifugus* BAR116 PhastCons element itself, and not the flanking sequence within the ChIP-seq peak, is essential for limb enhancer activity at this time point.

**Fig 5.**
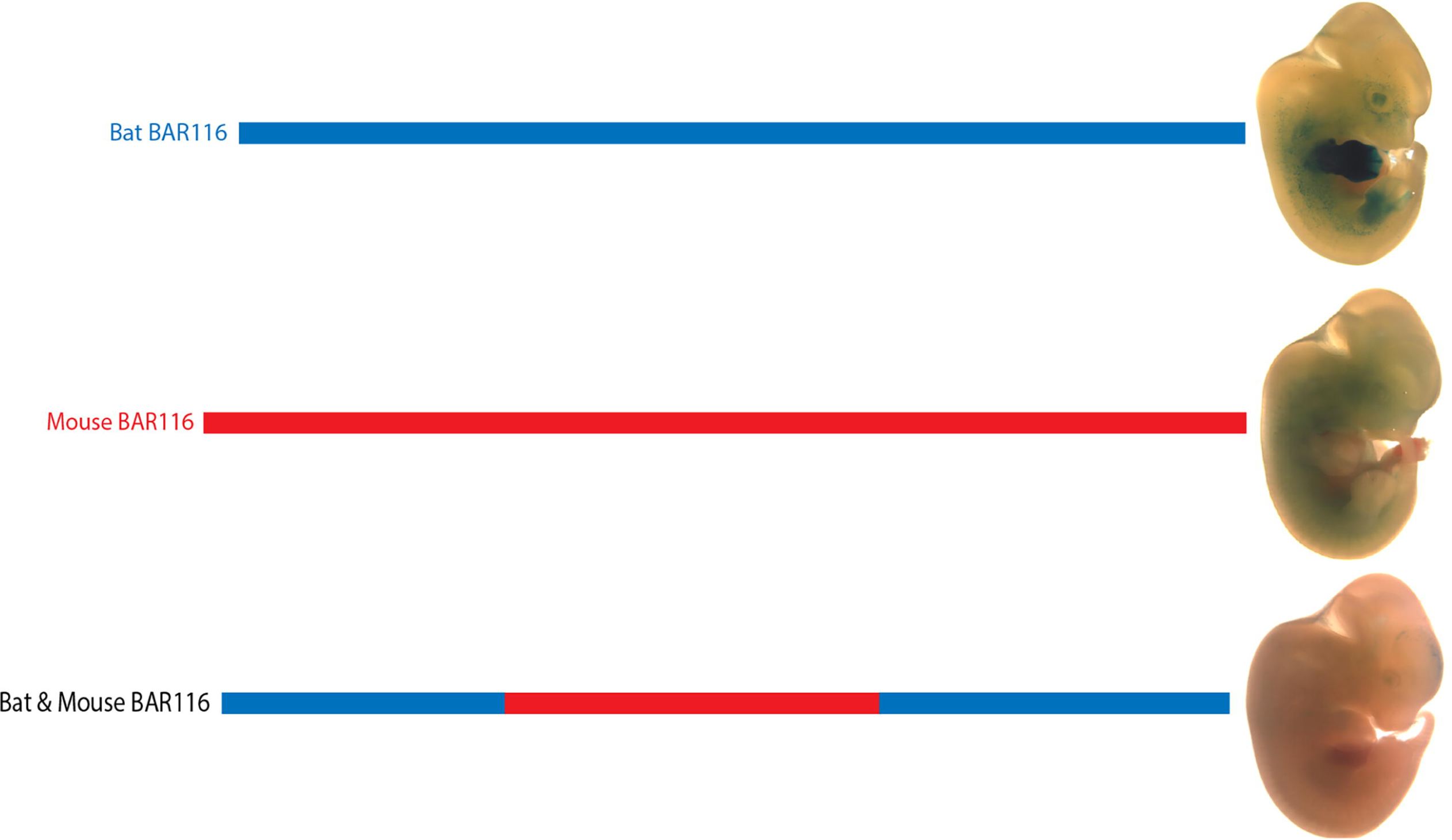
The bat-mouse BAR116 composite sequence displays a loss of tissue specificity in the limbs. The *M. lucifugus* sequence (blue) was replaced with the mouse (red) PhastCons sequence encompassing the BAR element. While the *M.lucifugus* sequence drove consistent enhancer activity in the limbs at E12.5, both the mouse sequence and the synthesized bat-mouse BAR116 composite sequence failed to drive consistent enhancer activity, suggesting that the *M. lucifugus* BAR116 sequence is essential for limb enhancer expression.

### BAR116 shows similar FL/HL expression differences to bat *Hoxd10* and *Hoxd11* genes

The most robust difference in enhancer activity observed was for *M. lucifugus* BAR116, which showed strong FL expression for the bat sequence but was negative for the mouse sequence. We wanted to analyze whether the *M. lucifugus* BAR116 enhancer expression pattern recapitulates that of the bat *HoxD* gene expression. We carried out whole-mount *in situ* hybridization on *Hoxd10*, *Hoxd11*, *Hoxd12* and *Hoxd13*, important developmental genes expressed during both early limb bud outgrowth and later autopod development [66], in developing bat (*M. natalensis*) and mouse embryos. At CS15, we observed FL expression of H*oxd10* and *Hoxd11* in both proximal (zeugopod) and distal (autopod) domains, while *Hoxd12* and *Hoxd13* expression were mainly limited to a large distal domain within the autopod (Fig 6A-D). The expression of all of these genes was strongest in a distal domain encompassing digits II-IV. The HL patterning at CS15 had both proximal and distal domains of expression for H*oxd10* and *Hoxd11*, while *Hoxd12* and H*oxd13* were found in the autopod region (Fig 6A’-D’). In the HL, the expression of the *HoxD* genes we analyzed appeared uniform and fairly symmetrical across the distal edge of the autopod. At CS16, the matching stage for E12.5 in mice, the bat autopod expands, becoming highly asymmetrical and digit rays become apparent. In the bat FL the distal expression of *Hoxd10* and *Hoxd11* are indistinguishable from one another, with strong expression occurring in a ‘triangular’ domain between digits II–IV (Fig 6E-F). *Hoxd12* has a more expansive autopod expression, extending from the posterior region of digit II and throughout the posterior portion of the autopod, with expression being focused in the proximal half of the developing digit rays (Fig 6G). *Hoxd13* was found throughout the autopod, and was strongest in the regions around the developing digits and in the interdigital region between digits III-V (Fig 6H). Interestingly, at CS16, we observed that *HoxD* expression in the HL is lost in the distal portion for *Hoxd10* and *Hoxd11*, while being maintained in the region where the calcar develops (Fig 6E’ & F). This differential expression in the FL and HL was similar for *M. lucifugus* BAR116, whereby we observed a robust FL expression within the entire limb but reduced HL expression in transgenic mice at E12.5. Expression of *Hoxd12* and *Hoxd13 at CS16* was maintained in the bat HL (Fig 6G’ & H’). In CS17 FLs, when the digit rays extend, expression of all the *HoxD* genes examined becomes progressively restricted to the regions surrounding the digits and is reduced in the distal interdigital tissue. In the HL, expression of *Hoxd10* and *Hoxd11* is still absent, and is reduced in the calcar region (Fig 6I’ & J’). *Hoxd12* expression appears reduced and is only found surrounding the digit rays while *Hoxd13* is found throughout the autopod region (Fig 6K’ & L’). In summary, although the observed expression pattern for *Hoxd10* and *Hoxd11* did not fully recapitulate the *M. lucifugus* BAR116 limb enhancer expression pattern, we did observe lower HL expression for both these genes at an equivalent stage that matches the decreased activity of the *M. lucifugus* BAR116 enhancer.

**Fig 6.**
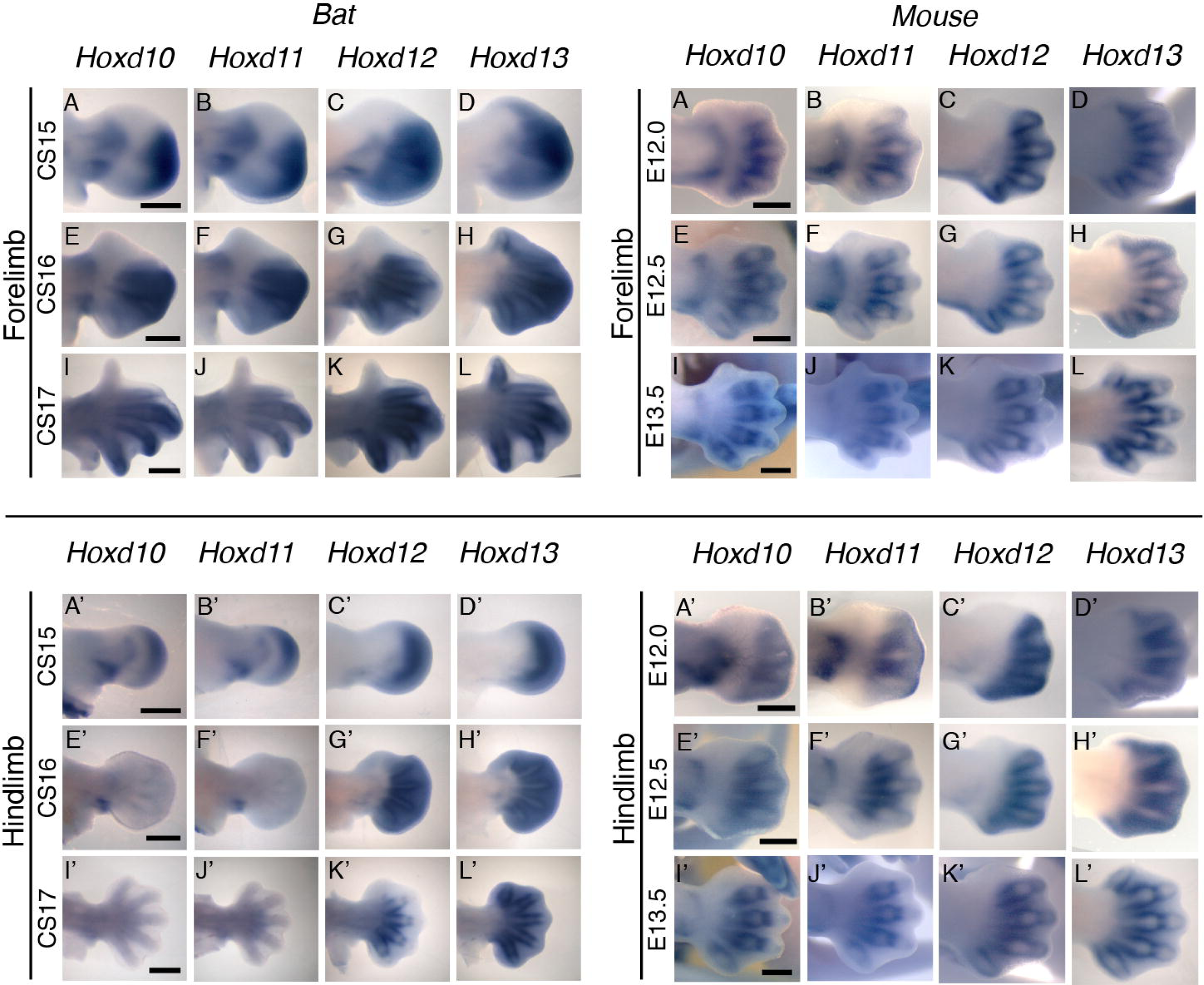
*HoxD* gene expression patterns in bats and mice. *Hoxd10-13* bat embryonic forelimb (A-L) and hindlimb (A’-L’) expression pattern at CS15-CS17 compared equivalently staged mouse (E12.0-E13.5) forelimb (M-X) and hindlimb (M’-X’) expression (scale bar represents 500 μm).

## Discussion

Using comparative genomics, developing limb ChIP-seq datasets and mouse enhancer assays, we characterized genomic regions in bat genomes that could play a role in mediating gene expression changes underlying the unique morphological development of the bat wing. We identified 166 BARs which showed a global enrichment for *Nr2c2*, *Zfp281*, *Zfp740*, *Zic2*/*3* and Egr1, and depletion for *Osr1*, *Osr2*, *Tgif1* and *Meis1* TFBS when comparing their mouse sequences to the inferred ancestral bat sequences (S2 Table). Analysis of five *M. lucifugus* BARs using a mouse transgenic assay showed all of them to have enhancer activity in the developing limb at E12.5. Examination of the mouse orthologous sequences for four of these BARs showed three to be differentially expressed compared to the *M. lucifugus* sequence, including BAR116 that showed strong FL versus HL expression, matching the differential expression pattern observed for *Hoxd10* and *Hoxd11* in developing bat limbs.

The *M. lucifugus* BAR4 sequence showed enhancer expression in the proximal FL [67], while the mouse orthologous sequence BAR4 was negative for enhancer activity at this time point (S1 and S2 Figs). BAR4 is near *Spry1*, a gene involved in skeletal and muscle development in mice that when overexpressed leads to chondrodysplasia, a skeletal disorder leading to arrested development [56,68]. Interestingly, 5 of our 166 identified BARs reside in the *Spry1* locus (Fig 2, S1 Table), suggesting that altered regulation of this gene could have been important for bat wing development. It is also worth noting that all five *Spry1* BARs are located in the same topological associating domain [TAD; [69]](Fig 2). The presence of multiple BARs in this region could implicate a co-operative and tightly coordinated regulation of *Spry1* during bat limb myogenesis and digit elongation via *Fgf* signaling [28,56]. The transcription factor *Plag1* could be differentially binding these enhancers and affecting the regulation of *Spry1* or other nearby genes.

Interestingly BAR61, the well-characterized *Shh* limb enhancer (ZRS), did not show differential enhancer activity between *M. lucifugus* and mice at E12.5. Both BAR61 sequences had similar enhancer activity in the ZPA of the autopod (S1 and S2 Figs.), however, both also seemed to have an expanded domain compared to the previously characterized E11.5 expression pattern [58]. These analyses do not exclude differences in expression pattern that may occur at later stages of development, or quantitative differences, which cannot be picked up through these transient transgenic assays.

BAR116 is of extreme interest, due to its telomeric location relative to the *HoxD* locus (Fig 2). The *HoxD* locus is conserved across vertebrates and has a critical spatiotemporal role in skeletal development [66]. *Hoxd10* and *Hoxd13* in particular are known to directly interact with *Shh* during limb outgrowth [70,71] and mutations in these genes lead to various limb malformations including synpolydactyly, split hand and foot, and distally located skeletal elements [66,72]. Several *cis* regulatory elements for the *HoxD* cluster have been shown to drive limb development and are conserved among vertebrates [10,73]. The *M. lucifugus* BAR116 portrayed enhancer activity in the FL mesenchyme but had reduced activity in HL (5/5 embryos). Its orthologous mouse sequence was negative for enhancer activity, despite showing evolutionary conservation in vertebrates. It is worth noting that a 700 bp partially overlapping region to mouse BAR116, termed CNS9 found in the *HoxD* telomeric region (Fig 2), was previously examined for enhancer activity in a lentiviral-mediated mouse transgenesis system and was also negative [74].

*M. lucifugus* BAR116 showed strong enhancer activity throughout the developing FL and weak expression in the proximal portion of the HL. Using whole-mount *in situ* hybridizations, we analyzed the expression patterns of *Hoxd10-13* at CS15-CS17 for *M. natalensis* and mice at matching developmental time points. Our results showed an overall similarity to previously characterized *Hoxd10*-*13* bat and mouse limb expression patterns [15,75]. The robust FL expression and weaker HL expression of BAR116 showed a similar trend to that of *Hoxd10* and *Hoxd11* (Fig 6). Interestingly, *Hoxd9* also showed a reduction in HL expression at CS16 [15], similar to the one we observed for *Hoxd10* and *Hoxd11*, suggesting that BAR116 could possibly be regulating these and other *HoxD* genes that are located 5’ to these genes. Additionally, by using 4C combined with *HoxD* telomeric deletion assays in mice, *Hoxd9*-*Hoxd11* were suggested to interact, during early phase of *HoxD* expression, with telomeric enhancers promoting forearm/arm development [74]. BAR116 (CNS9) lies within this telomeric region and is 850kb from the telomeric boundary of this topological domain suggesting that it could regulate *HoxD* genes. There are also two functional limb enhancers on both sides of it, CNS39 (60kb centromeric to CNS9; Fig 2) and CNS65 (229kb telomeric to CNS9; Fig 2), that are thought to interact with *HoxD* genes [74]. The differential enhancer activity we observed for *M. lucifugus* BAR116 compared to the negative enhancer activity of its mouse sequence and bat-mouse BAR116 composite sequence at E12.5, could imply that bats have acquired a novel enhancer function or a temporal specific chromatin conformation essential for forelimb morphology during autopod development in this locus.

Our data suggests that accelerated regions could be used to identify species-specific developmental enhancers that serve as critical determinants during morphoevolution as has been demonstrated previously [19,21,23]. In bats, an examination of the evolution of echolocation identified *Foxp2* as a major constituent of vocal and orofacial development pathways, finding conserved noncoding elements in close proximity of *Foxp2* that changed significantly in echolocating bats when compared to non-echolocating species [76]. Similarly, our study utilizes different species to test orthologous sequences, but focuses specifically on regions that are predicted to be developmental limb enhancers through ChIP-seq. There have been previous reports that analyzed bat-specific limb enhancers [12,75]. However, our study is the first to examine, in a genome-wide manner, putative limb enhancers in bats. Interestingly, the *Prx1* known bat limb enhancer whose replacement in the mouse led to longer forelimbs [12], was not identified in our analyses as a BAR element.

It is worth noting that our study also had many caveats. The mouse transgenic enhancer assay that we used is not quantitative and can be inconsistent due to differences in integration sites and transgene copy number. We also could not test sequences in bat embryos and so were limited to observing expression changes only in mice. For our *in situs*, due the scarcity of these embryos, we were only able to use *M. natalensis* embryos whereas in our mouse transgenic assays we used sequences from *M. lucifugus*, which could also lead to differences in expression patterns. Moreover, we only examined 5 of 166 BARs, which does not represent the majority of the BARs found in our pipeline. For our global TFBS analysis, we analyzed what we determined to be the ancestral bat sequence that could lead to us missing several TFBS changes. In addition, the TFBS matrixes we used were mainly human and mouse based and could differ in bats. In *M. lucifugus* BAR sequences we also noticed repetitive regions that were not present in the mouse BARs, which could explain the enrichment for specific TFBSs during our motifDiverge analysis. Despite these caveats, our study shows that the use of tissue specific ChIP-seq datasets combined with sequence acceleration can be an efficient means to identify sequences that are important in determining morphological changes between species.

## Materials and Methods

### Computational Analyses

To identify BARs, we employed a statistical phylogenetic test for accelerated nucleotide evolution in the common ancestor of all extant bats. This is an extension of a previously proposed likelihood ratio test for acceleration in a single species or clade [27]. This new ancestral lineage version of the likelihood ratio test is implemented in the PhyloP function (option –-branch) in the open source software package PHAST [77]. The input to PhyloP is a multiple sequence alignment for each genomic region to be tested for acceleration, plus a phylogenetic tree of the species in the alignment that is estimated from genome-wide data (in this case, four-fold degenerate sites).

To apply this statistical test to bat limb development, we first identified a collection of candidate enhancers for limb development genes by intersecting evolutionarily conserved elements with enhancer-associated histone modifications and transcription factor binding events measured in the developing mouse limbs (Fig 1). Specifically, we took the union of all peaks from two previously published ChIP-seq experiments targeting H3K27ac or p300 [17,26] and an H3K27ac dataset generated for this project. Next, we generated a set of vertebrate conserved elements that were agnostic to the rate of nucleotide substitutions in bats. We started with 60-way vertebrate multiple sequence alignments with mouse as the reference species (UCSC Genome Browser, mm10 assembly). We dropped the two bat genomes (*M. lucifigus* and *P. vampyrus*) from the alignments to ensure that high rates of nucleotide differences between the bats and other vertebrates would not prevent us from identifying conservation in other species. Finally, we ran the PhastCons program with default settings [25] on the resulting genome-wide alignments.

This analysis identified 4,384,943 conserved elements, many of which were less than 100 bp long and, thus, too short for statistical tests for acceleration [27]. However, we observed that many short elements frequently clustered together on the chromosome and that known functional elements (e.g., coding exons) were often tiled with multiple conserved elements separated by short gaps. Hence, we iteratively merged adjacent elements until the ratio of the distance between the elements merged over the total length of the region was less than or equal to 0.1. This merging algorithm was the result of empirical experiments aimed at producing one or a small number of merged elements per exon. We also experimented with adjusting the parameters of PhastCons to produce longer elements, but found that post-processing, by merging, recapitulated exons more effectively. Next, we intersected all merged regions greater than 100 bp with the ChIP-seq peaks and unmasked the *M. lucifigus* and *P. vampyrus* sequences from the multiple alignments. Regions with more than 50% missing sequence from either bat or more than 25% of nucleotides overlapping a coding exon were dropped to produce a collection of 20,057 candidate limb enhancers.

Prior to PhyloP analysis, we integrated sequences from two additional bat genomes into the candidate enhancer alignments. We obtained assembled contigs for two bats, *M. davidii* and *P. alecto*, that were sequenced to high coverage (100x)[13]. We used the BLAST algorithm to identify alignments of the mouse sequence from each candidate enhancer to contigs from *M. davidii* and *P. alecto* [78]. The single best hit with an e-value less than or equal to 0.01 was then blasted back to the mouse genome. If this produced a reciprocal best hit (i.e., the top scoring alignment to the mouse genome overlapped the original candidate enhancer sequence), we added the *M. davidii* or *P. alecto* sequence to the 60-way multiple alignment for that candidate enhancer. This produced alignments with between two and four bats present per enhancer. The two additional bat species were added to the phylogenetic tree corresponding to the 60-way alignments (UCSC Genome Browser) and their branch lengths were adjusted using their relationship to *M. lucifigus* and *P. vampyrus*. We then restricted our analysis to regions containing at least one bat.

Finally, we used PhyloP to test each candidate enhancer for accelerated nucleotide substitutions along the ancestral bat lineage. The resulting p-values were adjusted for multiple testing using a false discovery rate (FDR) controlling procedure [79,80]. We call all candidate enhancers with FDR < 5% Bat Accelerated Regions (BARs) (S1 Table). Their genomic distribution and sequence composition were analyzed using custom Python scripts. Significant associations with functions and phenotypes of nearby genes were identified using GREAT after lifting BARs over to mm9 coordinates [81]. We curated a list of limb-associated genes by exhaustively looking through the literature for evidence found in mouse or human and used resampling tests to assess associations between BARs and these genes compared to random sets of PhastCons elements.

### TFBS analyses

To look for TFBS differences, we manually curated a list of limb-associated TFs (S2 Table). BARs were analyzed for loss and gain of binding sites for each TF using motifDiverge [34]. We first compared the ancestral bat sequence to mouse. We used prequel to computationally infer the sequence of the common ancestor of extant bats using our multiple alignments [77]. We created the corresponding aligned mouse sequence from these alignments. We then called a TFBS a hit if its FDR exceeded a threshold of 0.01. We then used motifDiverge [34] to test if the total number of TFBS in the bat ancestor was significantly different than the number of TFBS in mouse for each TF in each individual BAR. We repeated these tests collectively over all BARs.

### ChIP-seq

ChIP was performed using the LowCell# ChIP kit (Diagenode) as previously described [82]. About 70,000 cells were pooled per IP and sonicated with a Covaris sonicator (S220 Focused-ultrasonicator, Covaris). Of the sheared chromatin, 30ul was used for each ChIP experiment with the antibody anti-acetyl histone H3 (Lys27) clone CMA309 (Milipore 05-1334). Following the manufacturer’s directions, each library was constructed using the Rubicon ThruPLEX library construction kit. Each library included 10 ul of ChIP material for a total of 14 cycles of amplification. Sequencing was carried out using an Illumina HiSeq and FASTQ files were aligned to the *Mus Musculus* genome (*mm9*) using Bowtie 0.12.8 [83]. A single base pair mismatch was permitted and reads with multiple alignments were discarded. The ChIP-seq library was sequenced to a depth of 168M total reads with 137M aligning uniquely. The input sample was sequenced to a depth of 111M reads total and 81M aligning uniquely. In each case, approximately 18% of sequences failed to align. We sorted and indexed the alignments using SAMtools 0.1.18 [84] and then converted to BED files with the bam2bed utility, a part of bedtools 2.17.0 [85]. To identify enriched H3K27ac islands in the limb samples, the peak-finding tool SICER 1.1 [86] was used.

### Mouse transgenic enhancer assays

PCR was carried out either on *M. lucifugus* or M. musculus DNA using primers that were designed to amplify candidate enhancer peak sequences with additional 100–500 bp outside of predicted regions (S1 Table). The bat-mouse BAR116 composite sequence was synthesized (Biomatik) and sequence validated. PCR products and the synthetic sequence were cloned into the Hsp68-LacZ vector [62] and sequence verified. All transgenic mice were generated by Cyagen Biosciences using standard procedures [87], and harvested and stained for LacZ expression at E12.5 as previously described [88]. Pictures were obtained using an M165FC stereo microscope and a DFC500 12-megapixel camera (Leica). To be designated as an enhancer, we required consistent spatial expression patterns present in at least two embryos. All animal work was approved by the UCSF Institutional Animal Care and Use Committee.

### Bat embryo collection

Ethical approval to collect bats was given by the University of Cape Town, Faculty of Science Animal Experimentation Committee (2006/V4/DJ, 2008/V16/DJ and 2012) with permission to sample granted by the Western Cape Nature Conservation Board (AAA004-00030-0035 (2006), (2008) and (2012). *M. natalensis* embryos were collected from the De Hoop Nature Reserve, Western Cape Province, South Africa in 2008 and 2012 and were harvested, staged and stored as described previously [63,65,89]. Whole-mount *in situ* hybridization was performed on wild-type Parkes wild-type strain (PKS) mouse embryos (Animal Ethics Committee application number: 006/040).

### Bat and Mouse whole mount in situ hybridization

Bat specific *in situ* probes were generated using primers that were designed to the *M. lucifugus* genomic sequence (Myoluc 2.0) and synthesized by Source Bioscience. PCR was performed using cDNA (S3 Table) and amplicons were gel extracted using the Wizard^®^ SV Gel PCR Clean-up System (Promega). Purified products were cloned using the pGEM^®^-T Easy Vector System (Promega) in XL-Blue cells. Purified plasmids (Promega Pure Yield^™^ Plasmid Miniprep System) from positively selected colonies were sequenced by Source Bioscience to confirm insert identity. NCBI blast analyses confirmed sequence similarity to the expected bat transcripts (S3 Table). Mouse probes (mHoxd10, mHoxd11, mHoxd12, mHoxd13) were generated from previously published probe templates [90]. Plasmids were linearized by digestion; bHoxd10: *NcoI*, bHoxd11-13: *SphI*, mHoxd10: *EcoRI*, mHoxd11: *SalI*, mHoxd12: *Bam*, mHoxd13: *pVUII* (Thermo Fisher Scientific) and purified to form an antisense probe template. DIG-labeled probes were generated using In Vitro Transcription (IVT). SP6 polymerase (Roche) was used for all bat probes and mHoxd10 and T7 polymerase (Roche) was used for mHoxd11-13 as per the manufacturer’s instructions. Reactions were DNAse treated using DNA-free^™^ (Ambion^®^, Life Technologies) and purified using a SigmaPrep^™^ Spin Column (Sigma-Aldrich). Embryos were sectioned along the sagittal plane to conserve samples and allow direct comparisons of different probes. Whole mount *in situ* hybridization was performed as described by [91] including Proteinase K digestion [92] and an overnight hybridization step at 70 °C followed by detection using NBT-BCIP.

## Accession numbers

ChIP-seq data has been made publically available through NCBI (ChIP-seq BioProject ID: PRJNA252737 as experiment ID SRX793524).

## Acknowledgements

We would like to thank Alisha Holloway and Dennis Kostka for lending us scripts to calculate significance of BAR and HAR clustering. We would also like to thank David Ray for the *M. lucifugus* DNA used in this work.

## Supporting Information

**S1 Fig. *M. lucifugus* BAR LacZ PCR positive embryos.** Insets showing higher magnification images of all embryos that had limb LacZ staining are shown next to the whole embryo.

**S2 Fig. Mouse BAR and bat-mouse BAR116 composite LacZ PCR positive embryos.** Insets showing higher magnification images of all embryos that had limb LacZ staining are shown next to the whole embryo.

**S1 Table. BARs identified through our computational pipeline.** BAR ID, ChIP-seq datasets where they originated from, ChIP-seq peak coordinates (mm10), PhastCons coordinates (mm10), PhyloP scores, p-Value (see computational methods), False Discovery Rates (FDR), and all genes within 1Mb upstream and downstream of BAR element are shown. The 38 BARs near limb-specific genes are highlighted in red font. The five mouse BARs that were found within gene deserts (gene desert defined by distance to a TSS that is greater than 500kb in either direction) are depicted with green highlighting. The *M. lucifugus* and mouse BARs chosen for enhancer assays are in bold and their primer sequences used for cloning are in the subsequent worksheet. To identify the nearest gene, we used LiftOver of the *M. lucifugus* sequence to mm9 coordinates.

**S2 Table. Limb-associated transcription factor binding site analysis.** The cloned sequences for *M. lucifugus* and mouse BARs were analysed for TFBS enrichment and depletion of known limb-associated TFs using motifDiverge [34,94]. We defined a TFBS if it exceeded an FDR of 0.01. TFBSs were tested for significant enrichment or depletion (Adjusted p-value < 0.05) in each cloned *M. lucifugus* BAR compared to its corresponding cloned mouse BAR. We also inferred the sequence of the common ancestor of the four extant bats for each of the 166 BARs using the prequel program in the PHAST package and compared these to their corresponding mouse sequences. We asked if TFBSs within these sequences were significantly enriched or depleted. All motifDiverge tests were corrected for multiple testing (to control the false discovery rate with the Benjamini-Hochberg method). Lists of all enriched/depleted TFs, limb-associated genes, and transcription factors are also provided.

**S3 Table. Bat *in situ* probe primers and GenBank numbers for the *HoxD* gene expression analysis.** Bat specific *in situ* hybridisation probes were generated using primers given. Here, we provide the amplicon lengths and GenBank identification numbers for Bat *Hoxd10-13* WISH probe sequences.

